# Hot-spots and their contribution to the self-assembly of the viral capsid: *in-silico* prediction and analysis

**DOI:** 10.1101/723023

**Authors:** Armando Díaz-Valle, José Marcos Falcón-González, Mauricio Carrillo-Tripp

**Affiliations:** Biomolecular Diversity Laboratory, Centro de Investigación y de Estudios Avanzados del Instituto Politécnico Nacional Unidad Monterrey, Vía del Conocimiento 201, Parque PIIT, C.P. 66600, Apodaca, Nuevo León, México; Unidad Profesional Interdisciplinaria de Ingeniería Campus Guanajuato, Instituto Politécnico Nacional, Av. Mineral de Valenciana No. 200, Col. Fraccionamiento Industrial Puerto Interior, C.P. 36275, Silao de la Victoria, Guanajuato, México

**Author notes:** Correspondence (M.C.-T.).

**Keywords:** free energy, structural conservation, functional dimer, protein–protein interaction, site-directed mutagenesis, binding free energy, molecular dynamics

## Abstract

In order to rationally design biopolymers that mimic biological functions, first, we need to elucidate the molecular mechanisms followed by nature. For example, the viral capsid is a macromolecular complex formed by self-assembled proteins which, in many cases, are biopolymers with an identical amino acid sequence. Specific protein-protein interactions drive the capsid self-assembly process, leading to several distinct protein interfaces. Following the hot-spot hypothesis, we propose a conservation-based methodology to identify those interface residues that are crucial elements on the self-assembly and thermodynamic stability of the capsid. We validate our predictions by computational free energy calculations using an atomic-scale molecular model of an icosahedral virus. Our results show that a single mutation in any of the hot-spots significantly perturbs the quaternary interaction, decreasing the absolute value of the binding free energy, without altering the tertiary structure. Our methodology can lead to a strategy to rationally modulate the capsid’s thermodynamic properties.

## 1. Introduction

The viral capsid is the archetypal molecular system to study the self-assembly of proteins. Capsid formation occurs rapidly and spontaneously with a high degree of fidelity. They are an excellent model for studying protein-protein interaction mechanisms in homo-oligomers that form symmetric closed shells. The viral capsid is a macromolecular complex formed by self-assembled capsid proteins (CPs) which, in many cases, are biopolymers with an identical amino acid sequence. It is believed that the self-assembly process has encoded signals in the sequence and structure of the CPs which direct the formation of the final virus particle. If present, such molecular recognition signals are crucial elements for the initial nucleation and subsequent growth of the capsid. Accurate identification of these molecular signals, so-called “hot-spots”, will help to understand viral self-assembly, and probably, macromolecular assembly in general. However, the molecular mechanism followed by the CPs to form the capsid is not fully understood yet due to the intrinsic complexity of the protein-protein interfaces.

Several works have proposed different strategies to predict the location of hot-spots. Many of the available methods take advantage of the fact that the contributions of interface residues to the binding free energy are not homogeneous. Usually, an energy-based alanine scanning mutagenesis method is implemented for this purpose. Examples of this type of predictors are ROBBETA [1], FoldX [2], SpotOn [3], and iPPHOT [4]. Other approaches have opted for machine-learning-based methods [5–8], molecular-dynamics-based methods [9,10], or combining solvent accessibility and inter-residue potentials [11]. Despite all these efforts, there is still not a clear way nor strict rule to locate hot-spots on a protein complex.

On a previous work, we reported the existence of a small set of interface residues for the icosahedral *Nodaviridae* virus family who are conserved not only in sequence but also in quaternary structure among all related members [12]. Furthermore, those structure-conserved interface residues were found to form non-random patterns around the capsid’s symmetry axes. That finding led us to believe other virus families might also present residues with those structural conservation characteristics. If true, it could be possible that those residues might have a crucial role in the self-assembly mechanisms due to their evolutionary relevance.

In this work, we propose an alternative methodology to locate hot-spots. We apply it to the *Bromoviridae* icosahedral virus family. Three steps can describe the general pipeline we followed: i) identify interface residues in each family member, ii) identify residues conserved in sequence, and iii) identify residues conserved in quaternary structure. The intersection of all these three sets are the hot-spots of that family. Furthermore, we also verified the hot-spot prediction by calculating the binding free energy change to the wild-type complex produced when mutating each one of the identified residues. Steered Molecular Dynamics (SMD) and Umbrella Sampling simulations were carried out to estimate the value of the binding free energy through the Potential of Mean Force (PMF) profile. Our findings show that our predicted hot-spots have a significant contribution to the complex formation, as opposed to other non-conserved interface residues.

## 2. Results

### 2.1. The composition of the CPs interfaces is not homogeneous

The Cowpea Chlorotic Mottle Virus (CCMV) belongs to the *Bromoviridae* family. We will use it as a representative member from this point on. The CCMV capsid protein residue composition is not homogeneous. An analysis of the CPs quaternary structure shows this holds for the interfaces too (Fig. 1). The structural regions found in a protein, namely, interface, core, and surface, grouped by their physicochemical nature are summarized in Table 1 for the case of the CCMV CP. A proportion of 40-40-20 percentage is found for residues on the interface, surface, and core, respectively. Half the total residues are non-polar, distributed in 20-20-10 proportion in the same order. Even though there is a couple of cysteines in the core, they are too far apart to form a disulfide bond. The small number of aromatic residues seem to be evenly distributed throughout the protein structure. However, a significantly larger number of charged residues are found in the interface with respect to any other structural region in this particular case.

**Table 1.** CCMV capsid protein residue composition by structural groups. Number of residues (%).

| Group | Non-Polar | Polar | Negative | Positive | Total | CYS | TRP |
| --- | --- | --- | --- | --- | --- | --- | --- |
| Interface | 33 (20) | 14 (9) | 12 (8) | 10 (6) | 69 (42) |  | W94 |
| Dimer<sup>a</sup> | 14 (9) | 5 (3) | 5 (3) | 3 (1) | 27 (16) |  | W94 |
| Core | 22 (13) | 8 (5) | 2 (1) | 1 (1) | 33 (20) | C59,C108 |  |
| Surf-Out | 31 (19) | 20(12) | 2 (1) | 2 (1) | 55 (34) |  | W55 |
| Surf-In | 4 (2) | 2 (1) | 0 (0) | 1 (1) | 7 (4) |  | W47 |
| Total | 90 (54) | 44(27) | 16(10) | 14 (9) | 164(100) |  |  |

**Figure 1.**
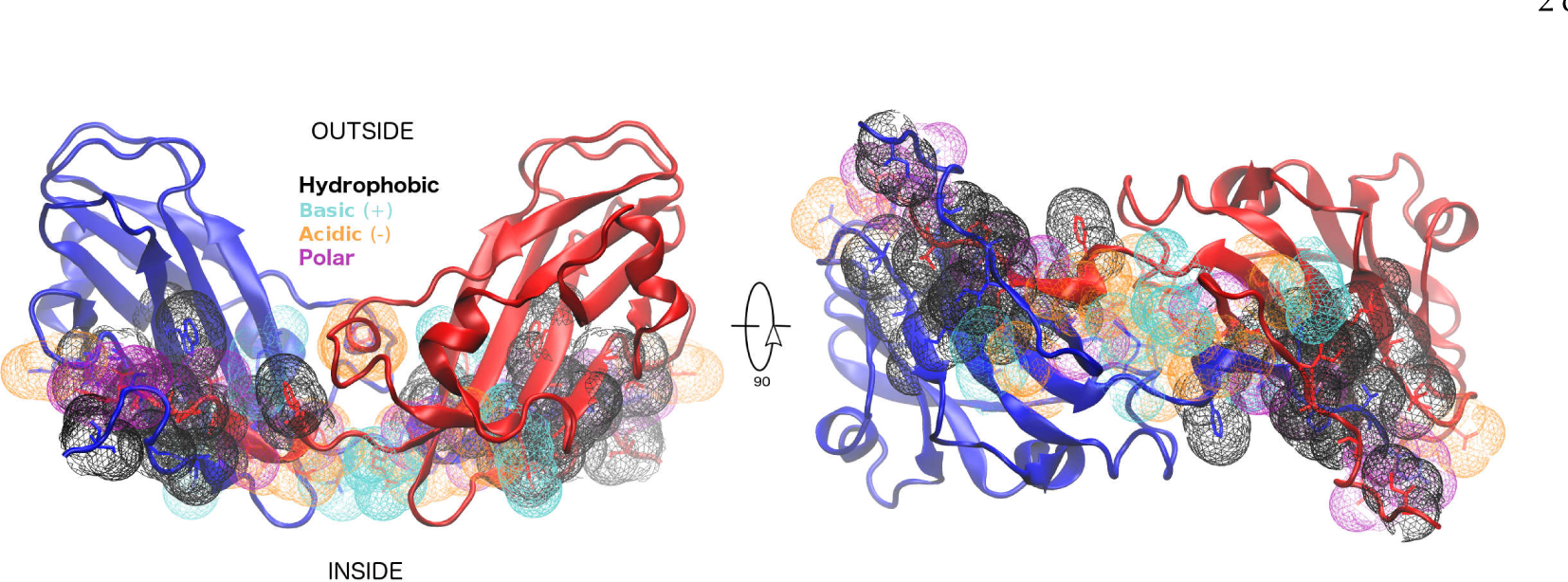
CCMV 2-fold related homodimer. Front view of the complex of subunit A2 (blue) and subunit B1 (red) in cartoon representation. The outside of the capsid is on top, the interior in the bottom. The dimer interface residues are shown in mesh representation, color-coded to represent the individual physicochemical property (hydrophobic, polar, basic, or acid). A 90-degree rotation has been applied on the right to appreciate the interface better.

Other physical properties have been used to characterize a protein. Some of these are the hydrophobicity (H) [13], the solvent accessible surface area (SASA) [14], the association energy (AEne) [15], the solvation energy (SolvEne) [16], and the buried surface area (BSA) [14]. We show a summary of these properties for the structural regions of the CCMV CP in Table 2 as reported on VIPERdb’s contact tables [17] (entry ID 1cwp). The total hydrophobicity by structural groups was estimated as the sum of the hydrophobicity index value of all residues in each group. As expected, the core and interface regions present large and equal values of hydrophobicity. Nonetheless, the surface exposed to the solvent also presents a large hydrophobicity. On the other hand, the surface in contact with the nucleic acids is rather hydrophilic. In terms of surface areas, the core region presents low values both in BSA and SASA. The outside surface presents low BSA but high SASA. The interface region has large values both in BSA and SASA. This observation could be related to the local flexibility of each region [18]. It is not surprising that most of the AEne come from the interface region.

**Table 2.** CCMV capsid protein physical-chemical properties by structural groups. Hydrophobicity (Mehler scale), Solvent Accesible Surface Area (Å^2^), Association Energy (kcal/mol), Solvation Energy (kcal/mol), and Buried Surface Area (Å^2^).

| Group | H | SASA | AEne | SolvEne | BSA |
| --- | --- | --- | --- | --- | --- |
| Interface | 18.5 | 2,382.3 | -147.1 | -51.4 | 6,351.9 |
| Dimer<sup>a</sup> | 8.6 | 864.2 | -87.1 | -34.4 | 3,707.0 |
| Core | 19.2 | 78.4 | -0.2 | -0.1 | 8.9 |
| Surf-Out | 14.6 | 2,853.5 | -3.4 | -0.8 | 171.1 |
| Surf-In | 2.2 | 604.3 | -2.5 | -1.2 | 118.4 |

### 2.2. There are six hot-spots in the Bromoviridae family

The multiple sequence alignment (MSA) of the *Bromoviridae* family members found in VIPERdb at the time of this writing, namely, Cowpea Chlorotic Mottle Virus (CCMV), Cucumber Mosaic Virus (CMV), Brome Mosaic Virus (BMV), and Tomato Aspermy Virus (TAV) is shown in Figure 2. The MSA was further corroborated by a tertiary structure alignment consensus made with all four proteins [19]. In general, the same capsid protein fold is conserved in all four family members. However, sequence identity is very low, highlighting a possible evolutionary structural convergence, as we had previously noted [20]. Less than 10% of the total residues are conserved in sequence. Half of those are found in the interface: P99, F120, Y159, H172, E176, R179, P188, and V189 (CCMV sequence numbering).

**Figure 2.**
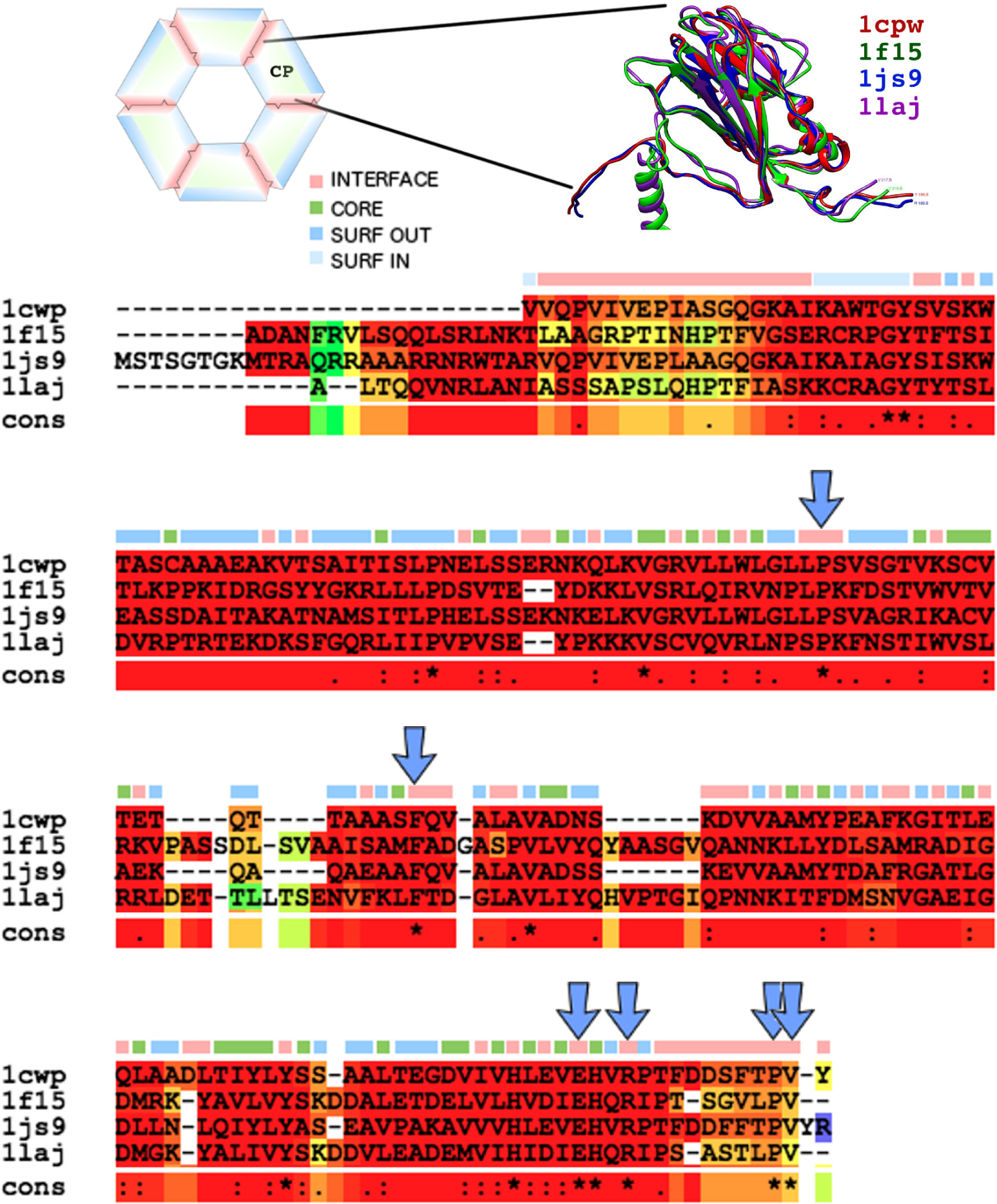
Multiple sequence alignment (MSA) of the *Bromoviridae* family members found in VIPERdb: Cowpea Chlorotic Mottle Virus (1cwp), Cucumber Mosaic Virus (1f15), Brome Mosaic Virus (1js9), and Tomato Aspermy Virus (1laj). The background color of the MSA represents tertiary structural similarity (low in green, medium in yellow, high in red). Top row, location of each residue in the capsids quaternary structure, mapping the sequence into four distinct groups (interface -red-, core -green-, outer surface -blue-, or inner surface -light blue-). Hot-spots are indicated with arrows. Inset) Multiple structure alignment of the capsid protein (CP) of 1cwp (red), 1f15 (green), 1js9 (blue), and 1laj (purple).

The multiple quaternary structure alignment of the capsid of the four viruses was achieved through the CapsidMap [21] methodology. A 2D depiction of the position of the residues in 3D is built by the projection on a plane. Then, a conversion from Cartesian to Spherical coordinates is made. The *φ*-*ψ* angle space is used to generate a map of the quaternary patterns formed by all the interface residues in a particular capsid. Individual CapsidMaps for the four viruses studied in this work are shown in Fig. 3. Conserved quaternary positions are readily identified when two or more CapsidMaps are compared. In the case of the *Bromoviridae* family, only six out of the eight interface sequence conserved residues are also conserved in quaternary structure.

**Figure 3.**
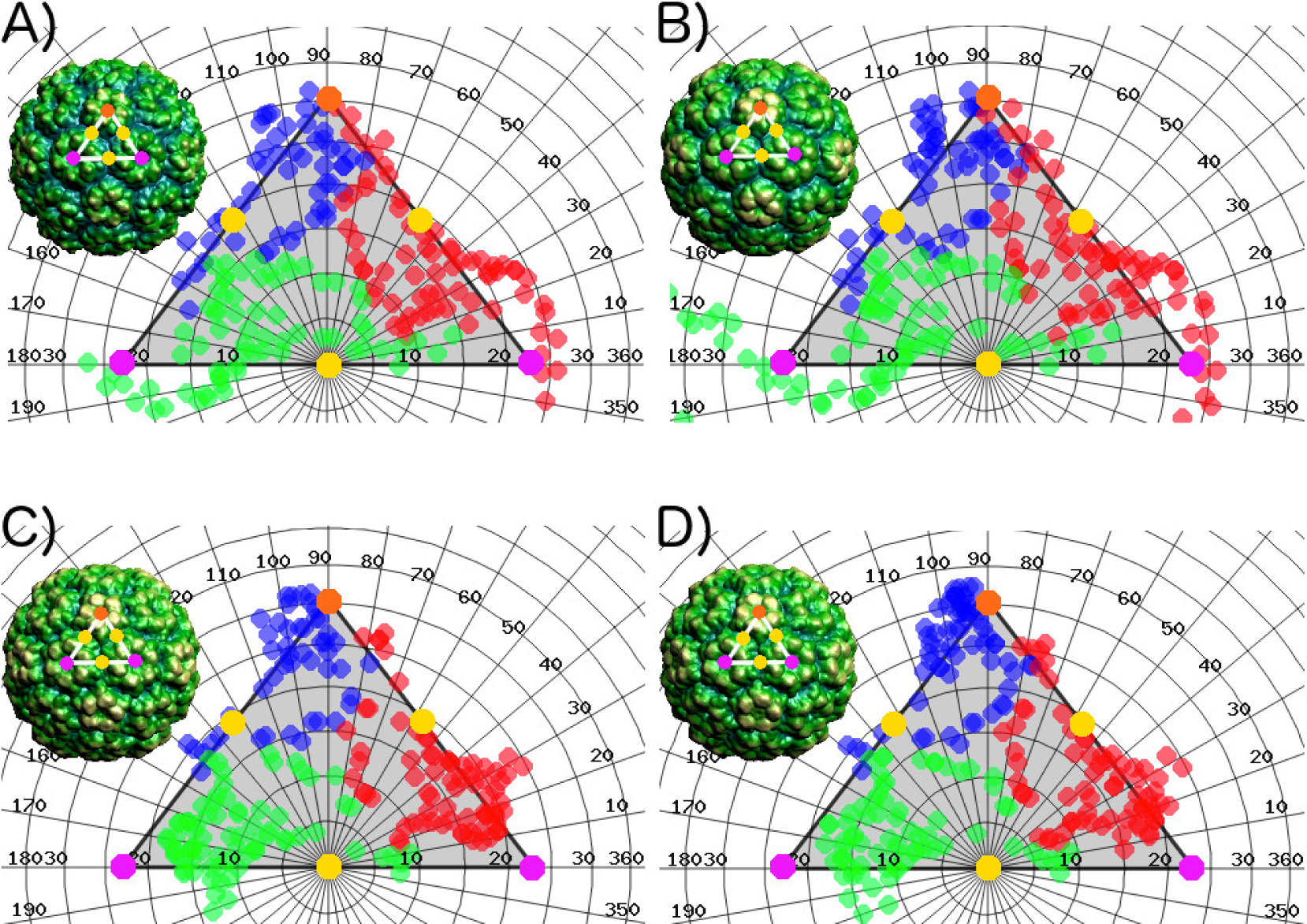
CapsidMaps of the interface residues of members of the *Bromoviridae* family, namely, (**a**) Cowpea Chlorotic Mottle Virus, (**b**) Brome Mosaic Virus, (**c**) Cucumber Mosaic Virus, and (**d**) Tomato Aspermy Virus. Each circle represents the location of an interface residue in *φ − ψ* space. The color used indicates the subunit to which the residue belongs (A in blue, B in red, and C in green). The icosahedral asymmetric unit used is indicated on the full capsid (insets), showing one 5-fold (orange), two 3-fold (magenta), and three 2-fold (yellow) symmetry axes. The 5-fold corresponds to *φ* = 90 *− ψ* = 30, and the opposite 2-fold corresponds to *φ* = *ψ* = 0.

Hence, the conservation criteria identify the set of residues P99, F120, E176, R179, P188, and V189 (CCMV sequence numbering) as hot-spots. Using the CCMV as a representative member, Figure 4 shows the location of these six residues on a CapsidMap. In order to assign each hot-spot to a specific fold-related interface, we used the VIPERdb contact tool (Virus Info Page-Annotations-Contact Tables-Which interfaces include a specific residue). Hot-spots P99 and F120 belong to the interfaces made around a 3-fold or a 5-fold, e. g., between subunits A1-A2, B1-C2, or B1-C6. On the other hand, hot-spots E176, R179, P188, and V189 belong to the interface made around a 2-fold, e. g., between subunits A1-B5, A2-B1, C1-C6, or C2-C9. The relationship between interfaces and symmetry folds and the location of the six hot-spots of the *Bromoviridae* family is illustrated in Figure 5. Any dimer related by the same type of symmetry fold is equivalent in the capsid quaternary structure. Given that there is ample evidence that the 2-fold related dimers are the first step in the capsid assembly process [22], we focused our efforts in the study of this particular type of interface in this work.

**Figure 4.**
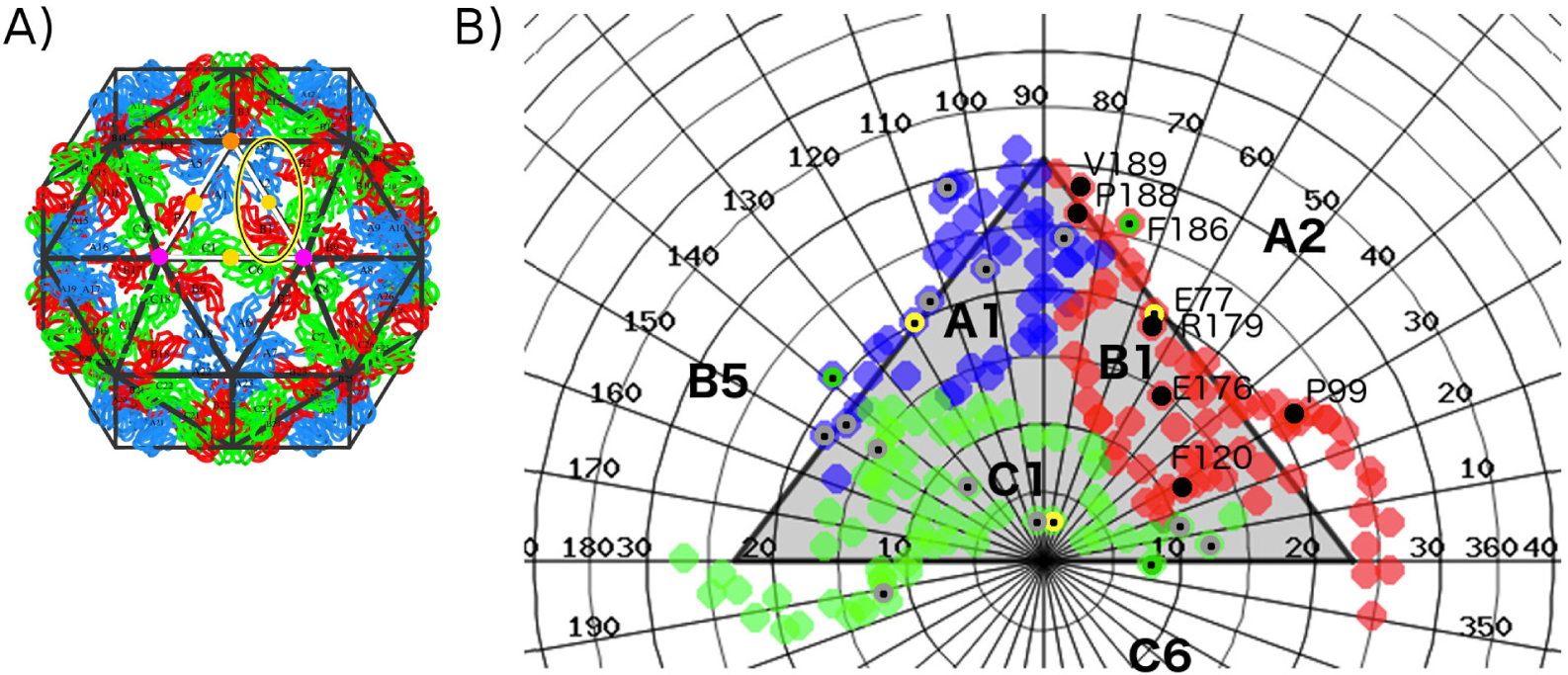
Multiple quaternary structure alignment of the *Bromoviridae* family members found in VIPERdb. The CCMV is used as the representative model. (**a**) Full capsid of the CCMV. The icosahedral asymmetric unit used in the analysis is indicated on white, showing one 5-fold (orange), two 3-fold (magenta), and three 2-fold (yellow) symmetry axes. The 5-fold corresponds to *φ* = 90 *− ψ* = 30, and the opposite 2-fold corresponds to *φ* = *ψ* = 0. All dimers related through a 2-fold axis are equivalent. One of them is highlighted, formed by the A2-B1 subunits. (**b**) CapsidMap of the CCMV interface residues, indicating the location of the hot-spots for subunit B1 (black dots), A1 and C1 (gray-black dots) in *φ − ψ* space. Location of the interface residues in the control group is also indicated (yellow and green dots).

**Figure 5.**
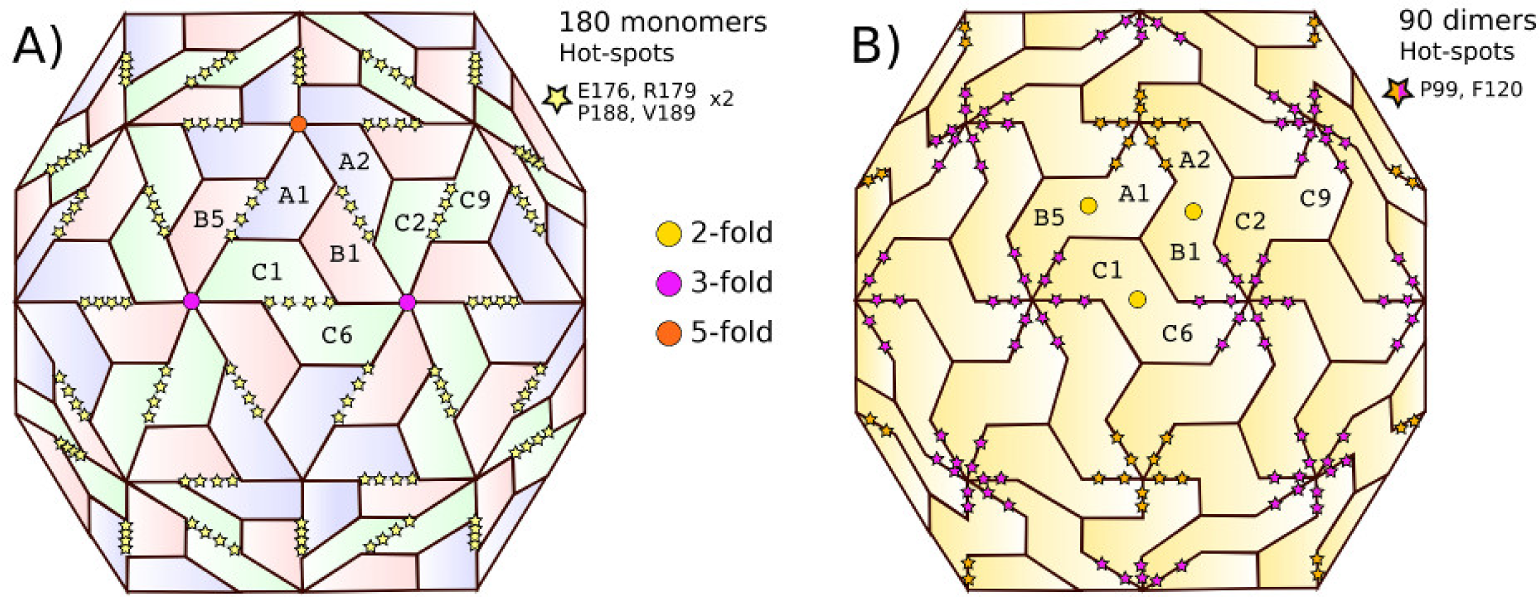
Mapping of the CCMV hot-spots on the full capsid. All hot-spots were found to be closely related to a given symmetry axis: (**a**) 2-fold (E176, R179, P188, V189), or (**b**) 3- and 5-fold (P99, F120). Symmetry axes are indicated for one of the 60 icosahedral asymmetric units. Only a few subunits are labeled.

We used the energy-based alanine scanning computational mutagenesis methods implemented in the ROBBETA, FoldX, SpotOn, and iPPHOT online tools to estimate the energy contribution of the interface residues of the 2-fold related A2-B1 dimer of CCMV. The SpotOn tool only reports a list of potential hot-spots (Fig. A1). The iPPHOT tool was not able to find any hot-spots on this protein complex (Fig. A2). ROBBETTA and FoldX both report the ∆∆G value by interface residue. A comparison between all hot-spot prediction results is presented in Figure 6. According to the energy-based alanine scanning analyses, there is an agreement on residue F186. In such a framework, this interface residue shows a significantly larger energy contribution than the rest of the interface residues, including the hot-spots predicted in this work. Given this observation, we included residue F186 in the following thermodynamic analysis.

**Figure 6.**
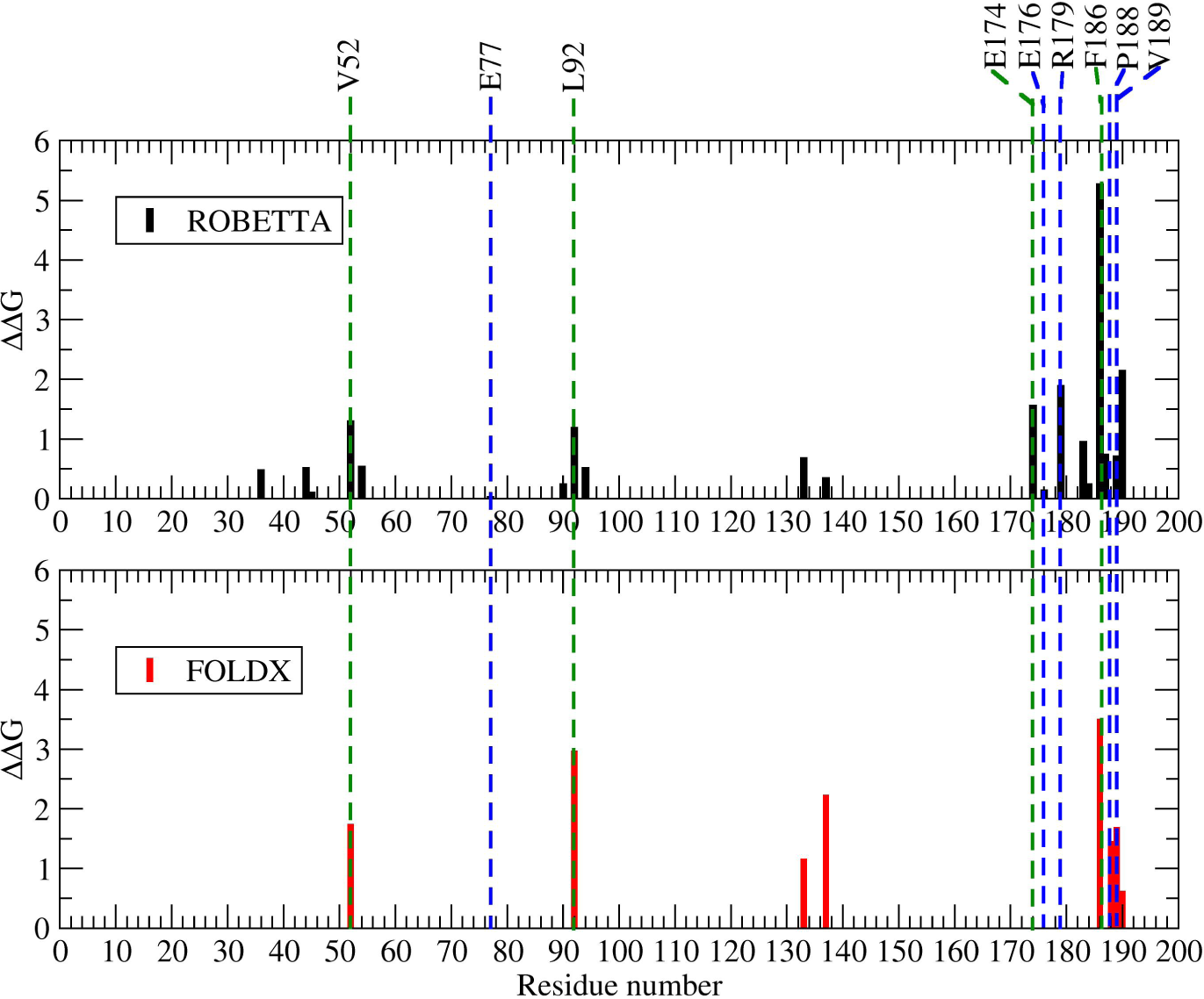
Computational energy-based alanine-scanning mutagenesis results from ROBBETA [1] (black), FoldX [2] (red), and hot-spot predictions from SpotOn [3] (green), and this work (blue) of a 2-fold related dimer interface. Interface residue E77 was randomly selected as an experimental control.

Furthermore, we randomly selected a non-conserved interface residue on the CCMV 2-fold related dimer as an experimental control (E77). Location in the quaternary structure of all the residues analyzed in this work is shown in Figure 7. A comparison of different characteristics shows that residue F186 stands out by having a low SASA and large BSA values, lower association and solvation energy values, as well as a significantly larger number of interactions with respect to the hot-spots predicted in this work (Table 3). These differences are probably the reason why all the energy-based prediction tools pointed it out.

**Table 3.** Hot-spots in the CCMV subunit dimer.

| Residue | SASA | AEne | SolvEne | BSA | NumInt |
| --- | --- | --- | --- | --- | --- |
| Proline P99<sup>a</sup> | 9.70 | -3.29 | -2.03 | 141.20 | 5 |
| Phenylalanine F120<sup>a</sup> | 11.10 | -0.73 | -0.35 | 34.73 | 1 |
| Glutamic Acid E176<sup>b</sup> | 0.00 | -0.16 | -0.10 | 6.74 | 2 |
| Arginine R179<sup>b</sup> | 64.50 | -2.80 | 0.32 | 138.64 | 6 |
| Proline P188<sup>b</sup> | 39.20 | -5.03 | -3.47 | 203.97 | 4 |
| Valine V189<sup>b</sup> | 28.70 | -4.48 | -2.92 | 187.98 | 7 |
| Glutamic Acid E77<sup>c</sup> | 65.50 | -2.72 | -0.54 | 112.78 | 2 |
| Phenylalanine F186<sup>c</sup> | 1.70 | -9.58 | -6.70 | 385.04 | 14 |

**Figure 7.**
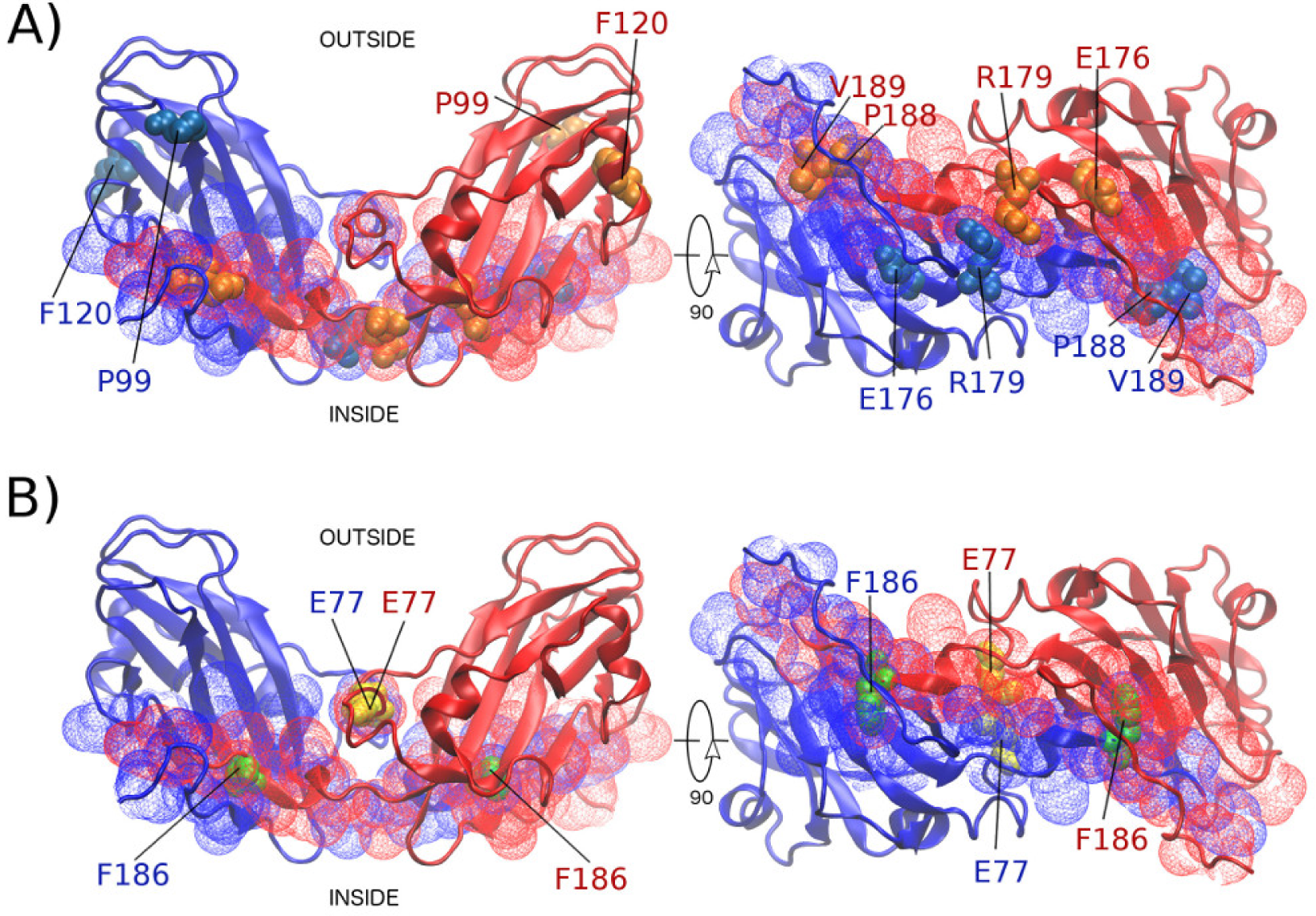
CCMV 2-fold-related dimer. (**a**) Location of the six hot-spots in the quaternary structure (space-fill representation), showing subunit A2 in blue and subunit B1 in red (cartoon representation). The dimer interface residues are shown in mesh representation. A 90-degree rotation has been applied on the right to appreciate the interface better. (**b**) Location of the interface residues in the control group (using the same representations as before).

### 2.3. Hot-spots significantly contribute to the protein complex binding

The MD trajectory data of all systems studied here can be accessed and visualized at the MDdb Science Gateway at http://www.md-db.org/id/690002. We mutated each hot-spot and control residue independently, producing seven variants. The rationale followed was to neutralize charges, change from non-polar to polar, or from big to small side chain, to disrupt all possible wild-type interactions. The final set was E176Q, R179Q, P188A, V189N, F186A, E77Q. The Umbrella methodology requires to sample conformations spaced along the reaction coordinate, which in this case is the distance between the center of mass (COM) of each subunit in the dimer. Hence, SMD trajectories were generated for the wild-type and the mutant set, starting from the homodimer complex and pulling the subunits until their COMs were 10 nm apart. MD simulations showed that none of the point-mutations disrupts the tertiary structure of the CCMV CP. Furthermore, except for mutant R179Q, there is no significant change with respect to the wild-type in the force needed to separate the protein complex (Fig. A3).

From the Umbrella sampling, the CCMV dimer potential of mean force (PMF) as a function of monomer separation for the wild-type, the point mutations in four hot-spots (E176, R179, P188, and V189), and the two residues in the control group (E77 and F186), was built with the WHAM algorithm (Figure 8). The computational cost was 17,500 CPUh, on average, for each one of the seven variants analyzed, totaling on an equivalent of 14 CPU years. As expected, the COM-COM distance in which the homodimer complex is formed corresponds to the global minimum in the PMF profile in all cases. It is clear that by pulling the two subunits 10 nm away was long enough to decrease their interaction energy to zero. This condition is sufficient to confidently assign the minimum point in the PMF profile as the free energy of dimerization in each case.

**Figure 8.**
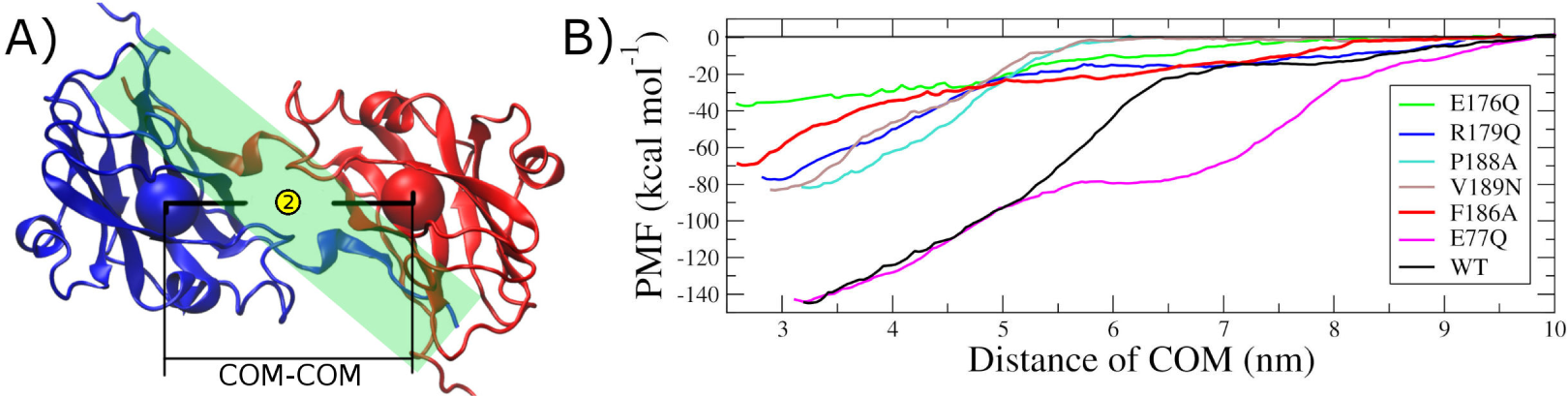
CCMV dimer potential of mean force (PMF). (**a**) The structure of a representative 2-fold related dimer (one subunit in blue, the other in red, cartoon representation). The reaction coordinate to generate the PMF profile was the distance between the center of mass (COM) of each subunit (spheres). The interface region is shaded. (**b**) PMF as a function of monomer separation for the wild-type and the point mutations in four hot-spots (E176, R179, P188, and V189) and the two residues in the control group (E77 and F186).

Hence, the change in the dimer interaction due to a point mutation relative to the wild-type, ∆∆G, is the difference found between them. Table 4 shows these differences for all the variants studied here. The randomly chosen interface residue E77 (experimental control) produces a positive change of less than 1 kcal/mol when mutated. On the other end, we found hot-spot E176, which produces a positive change of 108 kcal/mol. This value is close to 75% of the free energy of dimerization. The other three hot-spots also produce a positive change, decreasing the stability of the dimer by about 45%. Interface residue F186, although not a structurally-conserved hot-spot, produces a positive change close to 50%.

**Table 4.**
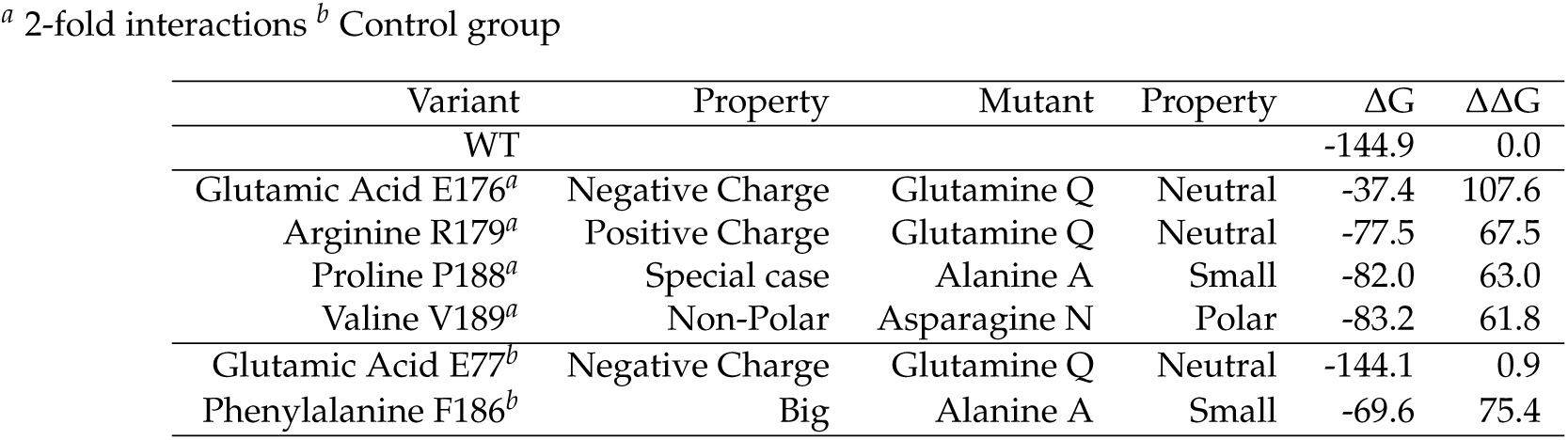
Free energy of dimerization, ∆G, and change in the dimer interaction due to a point mutation, ∆∆G, relative to the wild-type (WT) variant. All units in [kcal/mol].

| Variant | Property | Mutant | Property | $\Delta G$ | $\Delta\Delta G$ |
| --- | --- | --- | --- | --- | --- |
| WT |  |  |  | -144.9 | 0.0 |
| Glutamic Acid E176<sup>a</sup> | Negative Charge | Glutamine Q | Neutral | -37.4 | 107.6 |
| Arginine R179<sup>a</sup> | Positive Charge | Glutamine Q | Neutral | -77.5 | 67.5 |
| Proline P188<sup>a</sup> | Special case | Alanine A | Small | -82.0 | 63.0 |
| Valine V189<sup>a</sup> | Non-Polar | Asparagine N | Polar | -83.2 | 61.8 |
| Glutamic Acid E77<sup>b</sup> | Negative Charge | Glutamine Q | Neutral | -144.1 | 0.9 |
| Phenylalanine F186<sup>b</sup> | Big | Alanine A | Small | -69.6 | 75.4 |

## 3. Discussion

In this work, we present an alternative strategy to identify critical residues potentially relevant to the self-assembly of protein-protein complexes, in particular, icosahedral viral capsids, based only on structural conservation. Our findings provide evidence that a perturbation on any sequence-and-space conserved interface residue produces a significant decrease in the thermodynamic stability of the protein complex, as opposed to other interface residues. Seemingly, such hot-spots are important on the quaternary level but not necessarily so on the tertiary level. This statement implies that a mutation on any of those residues will not likely disrupt the protein fold but could prevent the complex from forming.

On a previous study, we applied the hot-spot prediction algorithm based on structural conservation to another virus family, namely, *Nodaviridae* [12]. Those results also showed the existence of six hot-spots. As in the case of the *Bromoviridae* family, their location in the capsid was not randomly dispersed throughout the protein-protein interfaces but forming patterns around the symmetry capsid folds. These findings are concomitant with the commonly accepted view of the capsid assembly kinetics. It has been shown that a nucleation seed needs to be formed in order to start the assembly process. At least in the case of the CCMV, this nucleation seed appears to be a pentamer of 2-fold related dimers (POD) [23].

Hot-spot identification is hard because there is no apparent correlation between residue type or interface composition with the way the complex is formed or the final relative orientation among subunits. Our hot-spot prediction methodology is straightforward, and even though it is not based on an energy calculation, here we have shown that the identified single residues do have a substantial energy contribution to the stability of the protein complex in comparison to non-conserved residues. The fact that an interface residue has been conserved in sequence and space in the quaternary structure during evolution can be explained if that particular residue plays a crucial role in the molecular mechanism of self-assembly, either to direct the process or to provide a stabilization anchoring point between the interacting proteins. Conservation seems to be a better search criterion than, for example, the number of contacts/interactions (E176 vs. F186, Table 3).

Energy-based alanine scanning is the standard way to search for hot-spots. When the energy function used is a rough approximation, it can be a fast method to estimate the energy contribution of all interface residues, one by one, to a protein complex. However, such approximation will not necessarily provide an accurate description of the molecular interaction. We used the method implemented in four available hot-spot prediction tools of this kind. Three of them identified residue F186 as the residue with a remarkable energy contribution to the complex. None of them picked out any of the six hot-spots identified by our methodology. Even more, our prediction based on conservation did not include residue F186 as a hot-spot. Nonetheless, all methodologies (energy or conservation-based) were in agreement concerning the interface residue E77 used as a control, which showed a contribution close to zero (Fig. 6).

Discrepancies found between our free energy estimates and those reported by the other methods are likely due to the different energy functions. For example, ROBETTA uses an effective energy function (averaging several contributions into one representation) and an isolated static molecular structure. Our mutagenesis scanning analysis (Umbrella sampling and PMF) employs a detailed description of all the bonded and non-bonded intra- and inter-molecular interactions, explicitly taking into account the contribution of the solvent, temperature, and pressure during a reasonable time, i. e., the whole system was thermodynamically equilibrated when calculating free energy.

Our methodology has limitations. First, the conservation-based prediction algorithm requires as many quaternary structures as possible, i. e., a single structure is not enough to detect interface residues conserved at the quaternary level. Second, if a thermodynamic confirmation is needed, the computation of the free energy is costly and requires the use of high-performance equipment to be viable. Nonetheless, this might not represent a problem nowadays with the increased access to supercomputing resources. All the computation packages needed in our methodology are free and open source.

Our mutagenesis free energy results show that the conservation-based hot-spot predictions have a large contribution to the formation and stability of the protein complex indeed. Results of *in-vitro* validation are reported in a companion article. In view of our findings, a similar study of the thermodynamic contribution of hot-spots P99 and F120 is in progress. These residues were excluded from this work because they are involved in protein interfaces other than the 2-fold related. Most likely, their role will be in the formation of intermediate states in the process of capsid assembly, e. g., PODs.

## 4. Materials and Methods

### 4.1. Multiple sequence alignment

The Geneious R7 software was used to carry out a multiple sequence alignment on a personal computer. The amino acid sequence of the members of the *Bromoviridae* family of which the crystallographic structure was available in the VIPERdb Science Gateway [17] were used.

### 4.2. Interface residues and quaternary structure alignment

We used the CapsidMaps [21] tool of VIPERdb to find the interface residues of members of the *Bromoviridae* family. The intra-family structural alignments were carried out using the method previously described [12]. The cured crystallographic structures of the viruses were queried to produce a CapsidMap of the interface residues. The *φ* and *ψ* coordinates of each residue were recorded and then compared between viruses. An overlap threshold of 3^*◦*^ in both angles was allowed to consider a conserved residue position in space.

### 4.3. Hot-spot in-silico mutations

The 3D structure of the wild-type dimer (WT) was obtained from the *Oligomer Generator* tool of VIPERdb. Indications were followed, and the necessary parameters were introduced to obtain a file in PDB format with the atomic coordinates of the selected dimer (A2, B1). All hydrogen atoms were added with the WHAT IF Web Interface. Histidine residues protonation state was set to physiological conditions (pH 7.0). Then, the *Mutator Plugin* tool of the Visual Molecular Dynamics software (VMD) [24] was used to generate one dimer with a point mutation for each one of the residues resulting from the structural conservation prediction. An additional point mutation was randomly selected from the A-B interface residues as an experimental control.

### 4.4. Alanine scanning mutagenesis

Comparison with other hot-spot prediction tools was made. We used the energy-based alanine scanning mutagenesis computational methods implemented in the ROBBETA [1], FoldX [2], SpotOn [3], and iPPHOT [4] (alignment created by ConSurf [25] using UNIREF90 and MAFFT) online tools. These are fast but coarse approaches for the prediction of energetically relevant amino acid residues in protein-protein interfaces. In all cases, the input was the 3D structure of the WT dimer in PDB format. The result was a list of residues predicted to significantly destabilize the interface when mutated to alanine, based on an approximated energy function.

### 4.5. Steered molecular dynamics

To accurately calculate the dimers binding free energy, a previously reported computational method using molecular dynamics was implemented [26]. We followed the same procedure for all the dimer variants produced in the previous section. The CHARMM27 all-atom force-field (FF) plus CMAP for proteins was used to describe molecular interactions [27]. This FF was chosen because it was shown that substantial deviations from experimental backbone root-mean-square fluctuations and N-H NMR order parameters obtained in the MD trajectories are eliminated by the CMAP correction, hence improving dynamical and structural properties of proteins. The dimers were solvated with liquid water using the TIP3P potential function [28].

All simulations were performed using the GROMACS 4.5.5 suite [29]. The initial structure of a dimer was placed inside a rectangular box. The dimensions of the simulation box were chosen such that the minimum distance between any atom of the peptide and the walls were no less than 1.0 nm and to provide sufficient space for the pulling to take place along the x-axis. The empty volume was filled with water molecules. Also, Na^+^ and Cl^*−*^ ions were added in such a proportion as to neutralize the overall charge of the system and to obtain a final salt concentration of 100 mM. Bad contacts between any two atoms were removed by energy minimization of the whole system using the steepest descent algorithm with a force tolerance of 100.0 kJ/mol/nm and a 0.01 step size. Isochoric-isothermal (NVT) equilibration of solvent molecules to 300 K was performed for 100 ps with the proteins heavy atoms being restrained by a harmonic potential with a force constant of 1000.0 kJ/mol/nm. A subsequent isobaric-isothermal (NPT) equilibration to adjust the system density was performed for another 100 ps at 1 bar.

Further structural equilibration simulations were carried out in the NPT ensemble at a temperature of 300 K and a pressure of 1 bar for 1 ns removing all position restraints. The temperature was maintained using the V-rescale thermostat with a coupling time constant of 0.1 ps. The dimer and solvent molecules were coupled to separate thermostats to avoid the hot solvent-cold solute issue. The pressure was regulated using the isotropic Parrinello-Rahman barostat with a coupling time of 2.0 ps and compressibility of 4.5×10^*−*5^ bar^*−*1^. Bonds involving hydrogen were constrained to their equilibrium values using the LINCS algorithm. The non-bonded interactions (Lennard-Jones and electrostatic) were truncated at 1.0 nm. Long-range electrostatic interactions beyond the cut-off distance were calculated using the particle mesh Ewald (PME) method with a Fourier spacing of 0.16 nm and a cubic interpolation of order 4. A long-range analytic dispersion correction was also applied to both energy and pressure to account for the truncation of the Lennard-Jones interaction. The time-dependent dynamics of the system was evolved using the leap-frog integrator with a time step of 2 fs.

After full equilibration, position restraints were set again for subunit A only, hence using it as an immobile reference for the pulling simulations. For each of the dimer variants, subunit B was pulled away from subunit A along the x-axis over 2000 ps, using a spring constant of 2000 kJ mol^*−*1^ nm^*−*2^ and a pull rate of 0.0035 nm ps^*−*1^ (0.035 Å ps^*−*1^). A final center-of-mass (COM) distance of approximately 7 nm between subunits A and B was achieved.

### 4.6. Umbrella sampling

From the previous SMD trajectories, snapshots were taken to generate the starting configurations for the Umbrella Sampling windows [30]. An asymmetric distribution of windows was used, such that the spacing was between 1.5 nm and 2 nm COM separation. Such spacing resulted in 42 windows per dimer. For each window, 5 ns of MD was performed for a total simulation time of 210 ns *×* 7 dimers utilized for umbrella sampling. Analysis of results was performed with the Weighted Histogram Analysis Method (WHAM) [31] for the generation of the Potential of Mean Force (PMF) profile as a function of the reaction coordinate (COM separation).

## Author Contributions

Conceptualization, M.C.-T.; methodology, M.C.-T.; validation, A.D.-V., J.M.F.-G. and M.C.-T.; formal analysis, A.D.-V., J.M.F.-G. and M.C.-T.; investigation, A.D.-V., J.M.F.-G. and M.C.-T.; resources, M.C.-T.; data curation, M.C.-T.; writing–original draft preparation, M.C.-T.; writing–review and editing, M.C.-T.; visualization, A.D.-V., J.M.F.-G. and M.C.-T.; supervision, M.C.-T.; project administration, M.C.-T.; funding acquisition, M.C.-T.

## Acknowledgments

All molecular dynamics simulations involved in the mutagenesis analysis reported here were performed with the *bmdhpc* computing resources of the Biomolecular Diversity Lab (tripplab.com) at CINVESTAV Unidad Monterrey, México, and the computing center *Insurgente* (www.cimat.mx/en/node/996) at the Centro de Investigación en Matemáticas, Guanajuato, México, thanks to the facilities kindly provided by Dr. Salvador Botello-Rionda from the Computer Sciences Department.

## Abbreviations

The following abbreviations are used in this manuscript:

CCMV: Cowpea Chlorotic Mottle Virus
CP: capsid protein
WT: wild-type
COM: center-of-mass
MD: Molecular Dynamics
SMD: Steered Molecular Dynamics

## Appendix A Supplementary data

**Figure A1.**
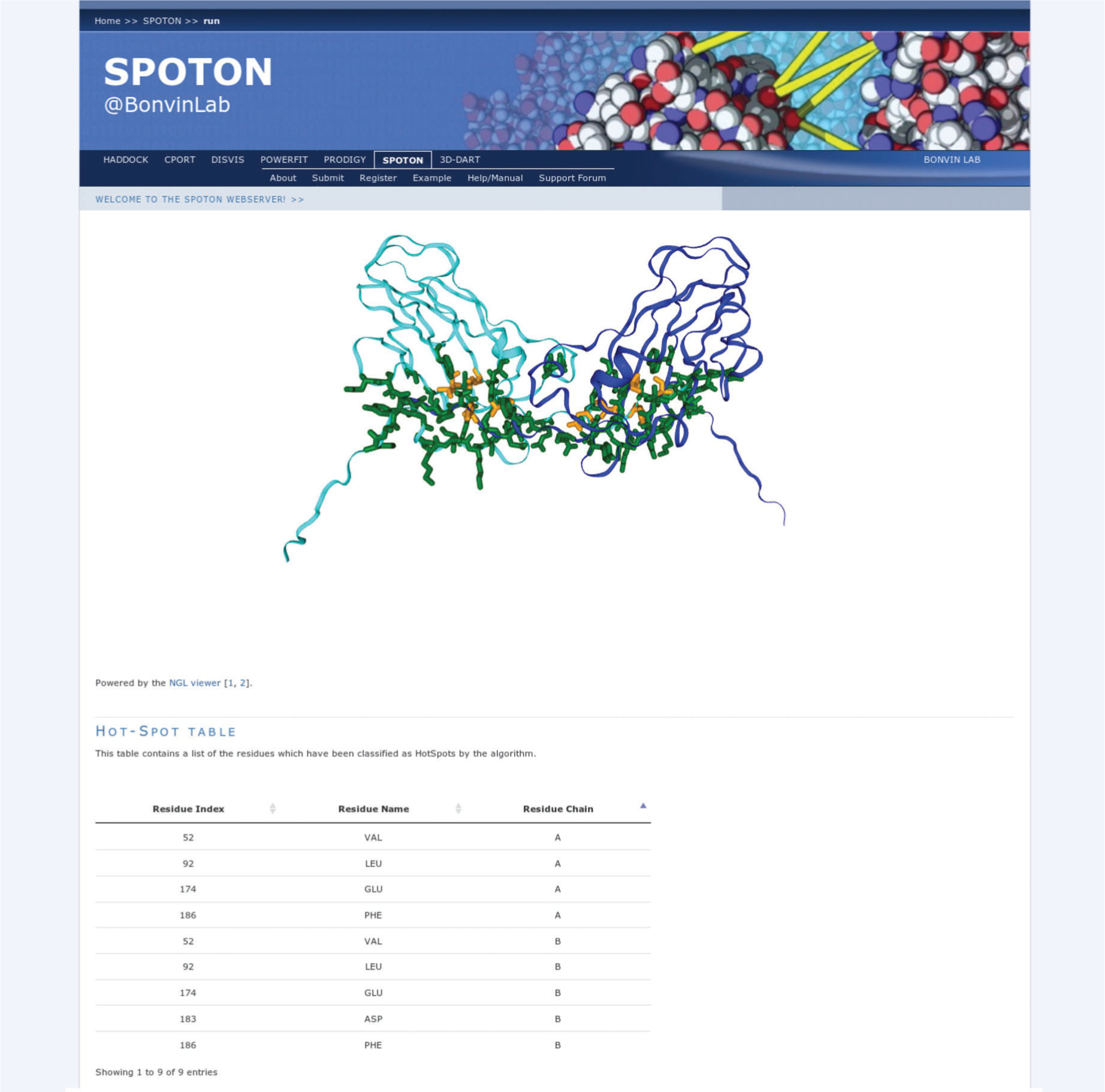
SpotOn [3] hot-spot predictions for the structure of a representative 2-fold related dimer.

**Figure A2.**
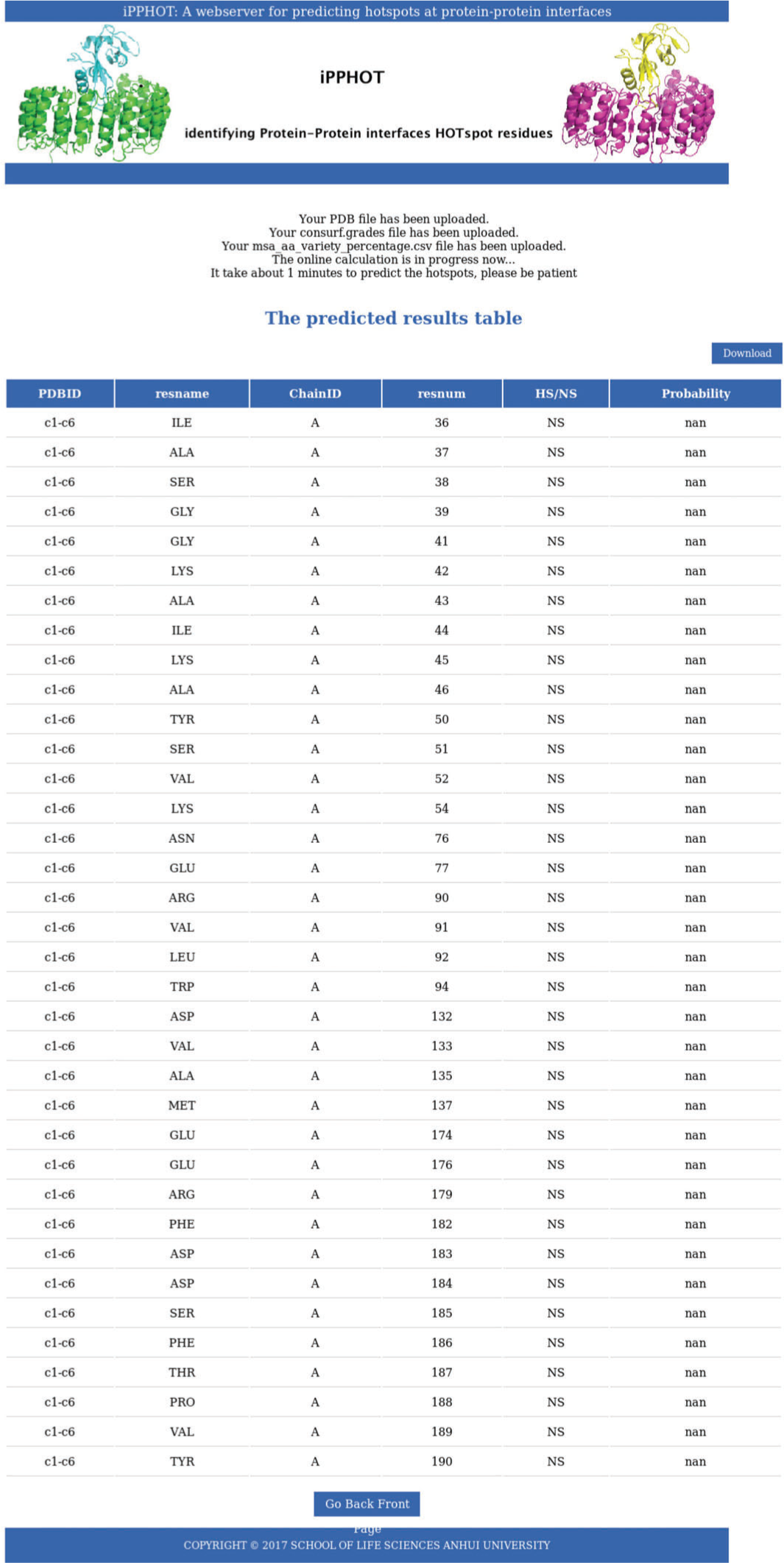
iPPHOT [4] hot-spot predictions for the structure of a representative 2-fold related dimer. All interface residues were predicted to be null-spots (NS).

**Figure A3.**
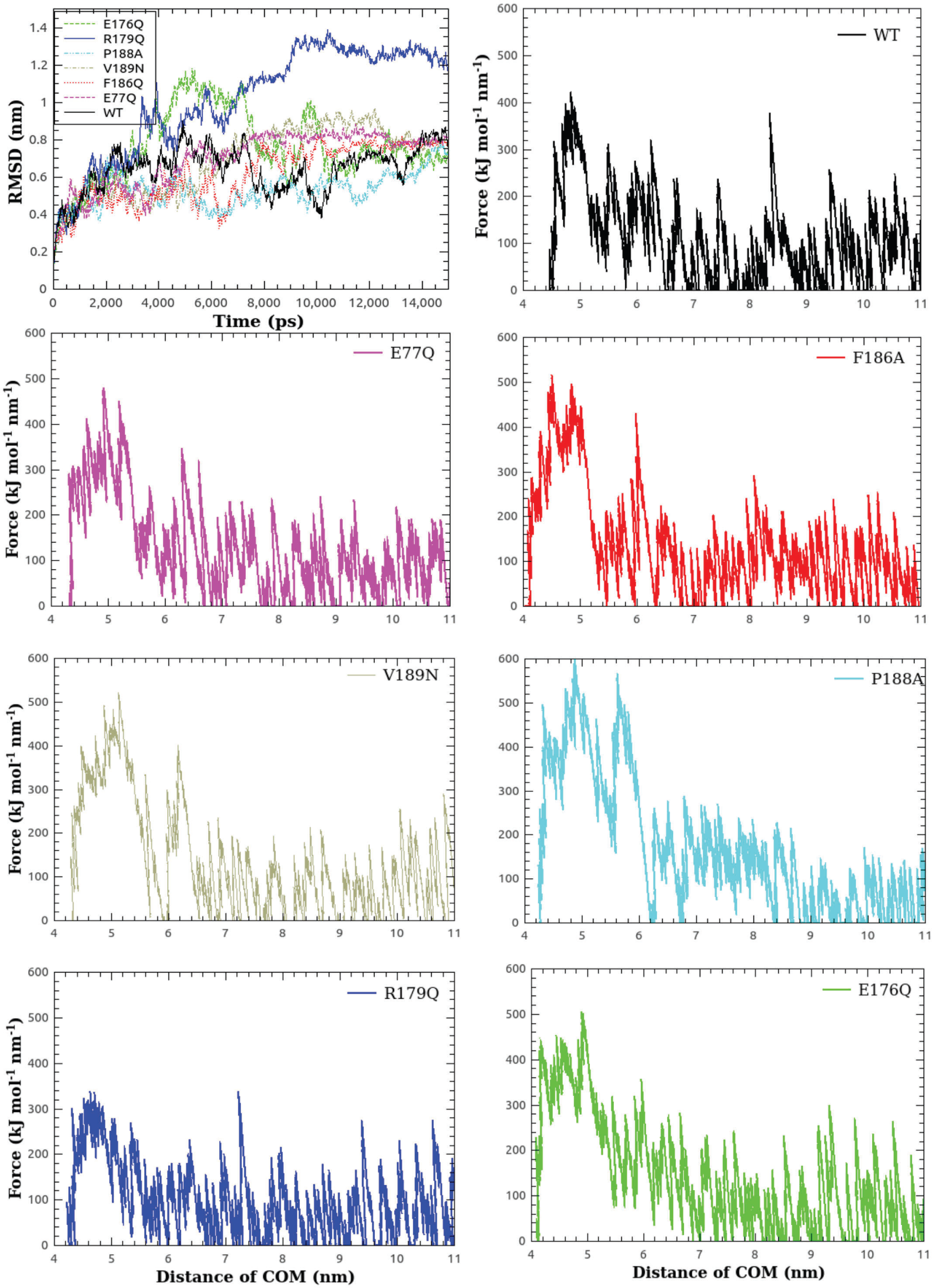
Monomer thermodynamic stabilization (RMSD vs. time), and 2-fold related dimer breaking force (F vs. COM-COM distance) for the seven CCMV CP variants studied in this work.

